# Reduced risk-taking behavior during frontal oscillatory theta band neurostimulation

**DOI:** 10.1101/2020.09.02.280594

**Authors:** Aline M. Dantas, Alexander T. Sack, Elisabeth Bruggen, Peiran Jiao, Teresa Schuhmann

## Abstract

**Background:** Most of our decisions involve a certain degree of risk regarding the outcomes of our choices. People vary in the way they make decisions, resulting in different levels of risk-taking behavior. These differences have been linked to prefrontal theta band activity. However, a direct functional relationship between prefrontal theta band activity and risk-taking has not yet been demonstrated.

**Objective:** We used noninvasive brain stimulation to test the functional relevance of prefrontal oscillatory theta activity for the regulatory control of risk-taking behavior.

**Methods:** In a within-subject experiment, 32 healthy participants received theta [4-8 Hertz (Hz)], gamma (30-50 Hz), and sham transcranial alternating current stimulation (tACS) over the left prefrontal cortex (lPFC). During stimulation, participants completed a task assessing their risk-taking behavior as well as response times and sensitivity to value and outcome probabilities. Electroencephalography (EEG) was recorded before and immediately after stimulation.

**Results:** Theta-band, but not gamma-band or sham, tACS led to a significant reduction in risk-taking behavior, indicating a frequency-specific effect of prefrontal brain stimulation on the modulation of risk-taking behavior. Moreover, theta-band stimulation led to increased response times and decreased sensitivity to reward values. EEG data analyses did not show an increase in power in the stimulated frequencies.

**Conclusion:** These findings provide direct empirical evidence for the effects of prefrontal theta-band stimulation on behavioral risk-taking regulation.

## Introduction

Human decision-making often includes a certain degree of risk regarding its outcomes and outcome probabilities. Take for example financial investments, driving above the speed limit or simply trying a new cuisine. In all of these situations, and many others, the outcomes of our decisions and actions cannot be predicted with absolute certainty. The decision-maker exhibits risk-taking behavior in these situations. Risk-taking is inevitable and may not only have (un)desired personal, but also social and economic impacts (1). Therefore, the regulatory control of risk-taking behavior is of utmost importance for human decision-making.

During decision-making under risk, a complex mechanism is at work. This mechanism codes and flexibly evaluates the context, outcome probabilities and previous information to define the optimal level of risk to be taken (2). Despite what is expected from pure rational models, risk-taking behavior is not consistent across different contexts and the most optimal decision is often rejected (3). These inconsistencies are likely a consequence of the complexity of the neural mechanisms involved in risk-taking behavior and the control thereof, which have been extensively explored by previous studies (e.g. 3–6). Namely, risk-taking behavior is the result of a complex interplay between emotional responses to possibilities of reward (limbic activity) and inhibition of such responses via activation of frontal control regions (7). Among these, the ventromedial prefrontal cortex (VMPFC) and dorsolateral prefrontal cortex (DLPFC) are critical areas, responsible for signaling the need of strategy adjustment and executive control, respectively (8). However, the exact mechanism underlying such signaling processes remains unclear.

Electroencephalography (EEG) studies have shown that participants with a higher theta power [4-8 Hertz (Hz)] in the right prefrontal cortex (rPFC) compared to the lPFC, i.e. a higher frontal theta-band asymmetry, displayed more risk-taking behavior during gambling tasks (9,10). These results are consistent with studies showing a positive correlation between error detection, cognitive control and increased right VMPFC theta power (11). Moreover, theta oscillations are involved in neural network communication when cognitive control is required (12). Prefrontal theta oscillations may therefore represent part of the signaling mechanism by which the VPMFC recruits the DLPFC, in case recruitment of regulatory mechanisms is needed because a risky context is detected. However, although these EEG studies indicate that theta oscillations are related to risk taking behavior, the functional behavioral relevance of this oscillatory pattern in the regulation of risk-taking has yet not been shown.

Noninvasive brain stimulation, such as transcranial alternating current stimulation (tACS), offers the possibility of inducing temporary oscillatory patterns in specific brain regions, by applying changing electric currents on the scalp transiently modulating brain activity. This allows to probe the relationship between frequency patterns and behavioral responses (13). To investigate the role of theta band frontal asymmetry in risk-taking behavior, Sela and colleagues (2012) applied theta-band tACS over right and left DLPFC while participants performed the Balloon Analog Risk Task. After left, but not right, DLPFC theta-band stimulation an increase in risk-taking behavior was found. This was not in line with prior EEG studies hypothesizing that right DLPFC stimulation increases risk-taking behavior, by increasing frontal asymmetry, while left DLPFC tACS reduces risk-taking behavior due to an increase in theta-band activity in the left hemisphere and consequent asymmetry reduction (6,9). Sela and colleagues (2010) speculated that their findings may be due to a disruption of interhemispheric balance in participant’s natural frontal asymmetry (14). The authors were not able to make conclusions about the frequency specificity of the stimulation, as no control frequencies were applied(15). Moreover, they opted for using the Balloon Analog Task to estimate risk. This task mostly measures impulsivity and evaluates uncertainty, rather than risk, which is a different economic construct (16), since the probabilities are not explicit to participants.

The present study aims at investigating the functional relationship between frontal theta-band oscillations and risk-taking behavior. Although previous studies (9,10) have shown a correlation between resting state frontal theta band asymmetry and risk-taking behavior, no direct causal relationship has thus far been shown. We therefore applied tACS to the left DLPFC in theta-band (6.5 Hz), and gamma-band (40 Hz) frequency as well as sham stimulation while participants performed a risk-taking task. We chose gamma-band tACS as a control frequency since it has not been linked to risk-taking behavior thus far. We also implemented a new behaviorally-controlled risk-taking protocol paired with financial incentives for more robust measures of risk-taking behavior.

To monitor possible power changes in the stimulated frequencies and to investigate their functional relationship with the behavioral results, we implemented EEG measurements before and immediately after the transcranial brain stimulation. We hypothesized that, compared to sham and gamma-band stimulation, theta-band stimulation to the left DLPFC decreases risk-taking behavior, confirming the central regulatory role of theta-frequencies on the electrophysiological mechanism underlying the modulation of risk-taking behavior (9,17).

## Material and methods

### Participants

Thirty-two healthy, right-handed students (16 female, mean age 23.8 years, range 18-31 years, SD= 3.45), participated in this study. All participants had normal or corrected-to-normal vision and gave written informed consent after being introduced to the experiment. They were screened for tACS safety, following the recommended procedures of Antal and colleagues (2017) (18), screening for e.g. skin diseases, implants, neurological disorders, pregnancy and medication.

The study was approved by the Ethics Review Committee Psychology and Neuroscience (ERCPN) of Maastricht University, The Netherlands (ERCPN 188_07_02_2018). Participants were compensated, in the form of vouchers with monetary value, based on the choices they made and luck in the risk-taking task, and for participating in the experiment. The stimulation was well tolerated by 31 out of 32 participants. One participant reported skin redness in the area of the stimulation after participating in session 1 and therefore decided to stop participation in the experiment. The results of this participant were excluded from the analyses.

### Procedure

Each participant received theta-band (6.5 Hz), gamma-band (40 Hz) and sham tACS in three separate sessions. The sessions were separated by an average of 7 days (+/−1) to avoid carry-over effects. Fig 1 provides an overview of our procedure and experimental design, which will be explained in the following sections. The order of stimulation conditions (interventions) was randomized across participants.

**Fig 1.**
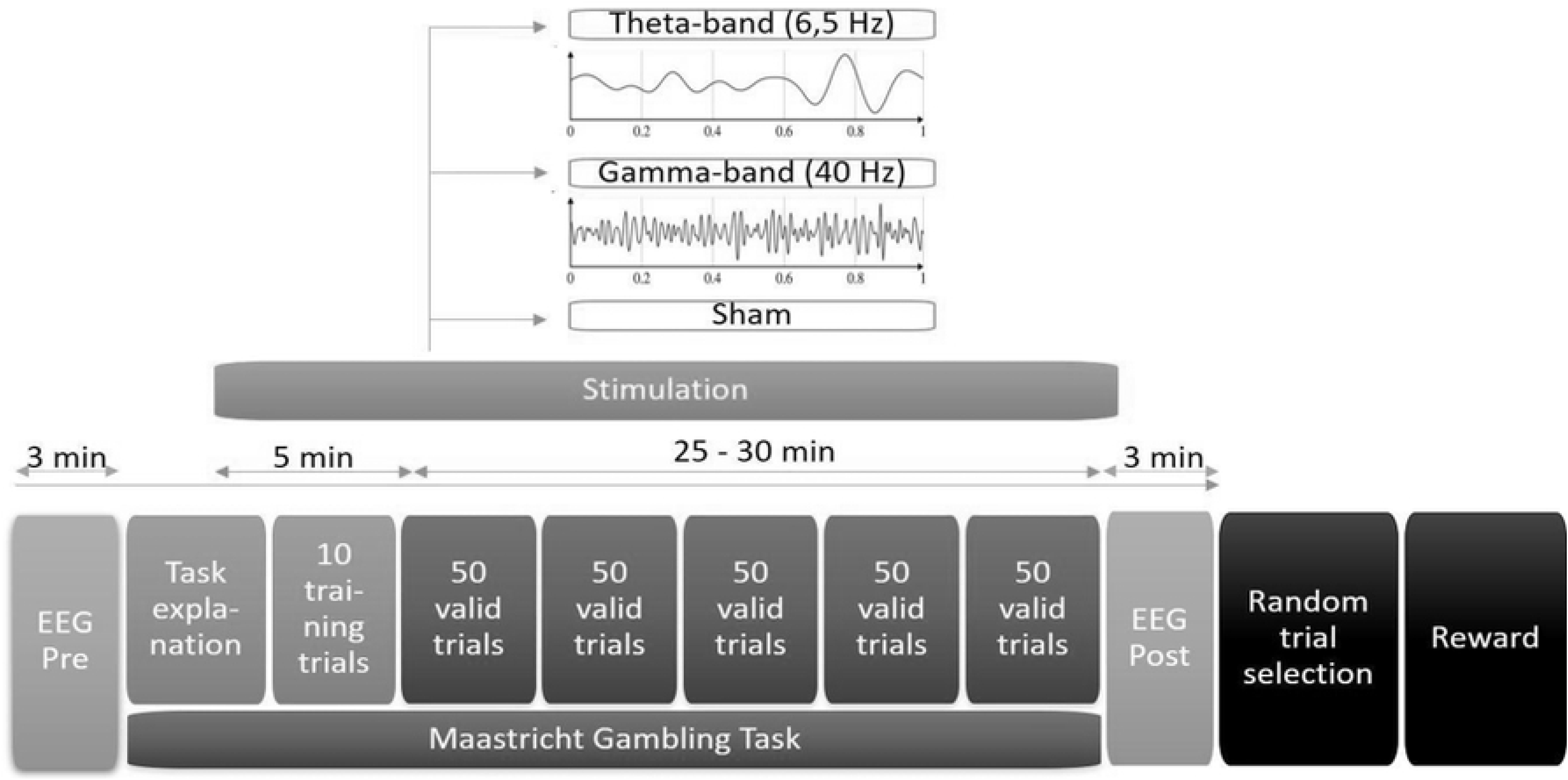
Experimental design. Experimental procedures, task presentation and experimental conditions.

Participants were invited to the laboratory where they reviewed the participant’s information, signed the safety pre-experimental check and the written consent and. In each session, participants were instructed about the experimental procedures and task and positioned at the workstation where the tACS and EEG electrodes were placed. EEG was measured before and after the stimulation. During tACS, participants had to perform a computerized gambling task named Maastricht Gambling Task.

Participants were informed that at the end of the session, one random trial of the task, which is composed by multiple trials (more details will follow in the corresponding session) would be selected for payment. They were asked to use an online random number generator to select the number of the trial that would be paid out. This was done in each of the three sessions. During the task, experimental currency was expressed as points. Every point earned in the selected trial was converted to € 0.10. All participants received also a participation fee of € 7.5 per hour (1.5 hour per session). The payments varied between € 33.75 and € 63.75 and were made only after the third session. All task details and payment rules were explained before the task (Fig1).

### Maastricht Gambling Task

A customized experimental protocol to elicit and access risk-taking behavior was developed based on the widely used “Risk task” (19) also known as Cambridge Gambling Task (CGT). The CGT is a valid measurement of risk-taking behavior (20,21), controls for impulsivity and was used in multiple studies using noninvasive brain stimulation (4,22–24). However, the CGT does not control for memory and wealth effects because the trials are not independent, meaning that participants carry gains and losses from the previous trials; and it is confounded by loss aversion, as participants could lose points during the task.

Therefore, we developed a revised protocol named the Maastricht Gambling Task (MGT). This computerized task presents six boxes (see Fig 2 for an example screen) to the participant, which can be either colored pink or blue. The number of pink boxes is randomized and ranges from 1 to 5, with the remaining boxes being blue. One of the colored boxes hides a token (represented by a yellow X) and the participant has to guess the color of the box that hides the token by choosing left (pink) or right (blue).

**Fig 2.**
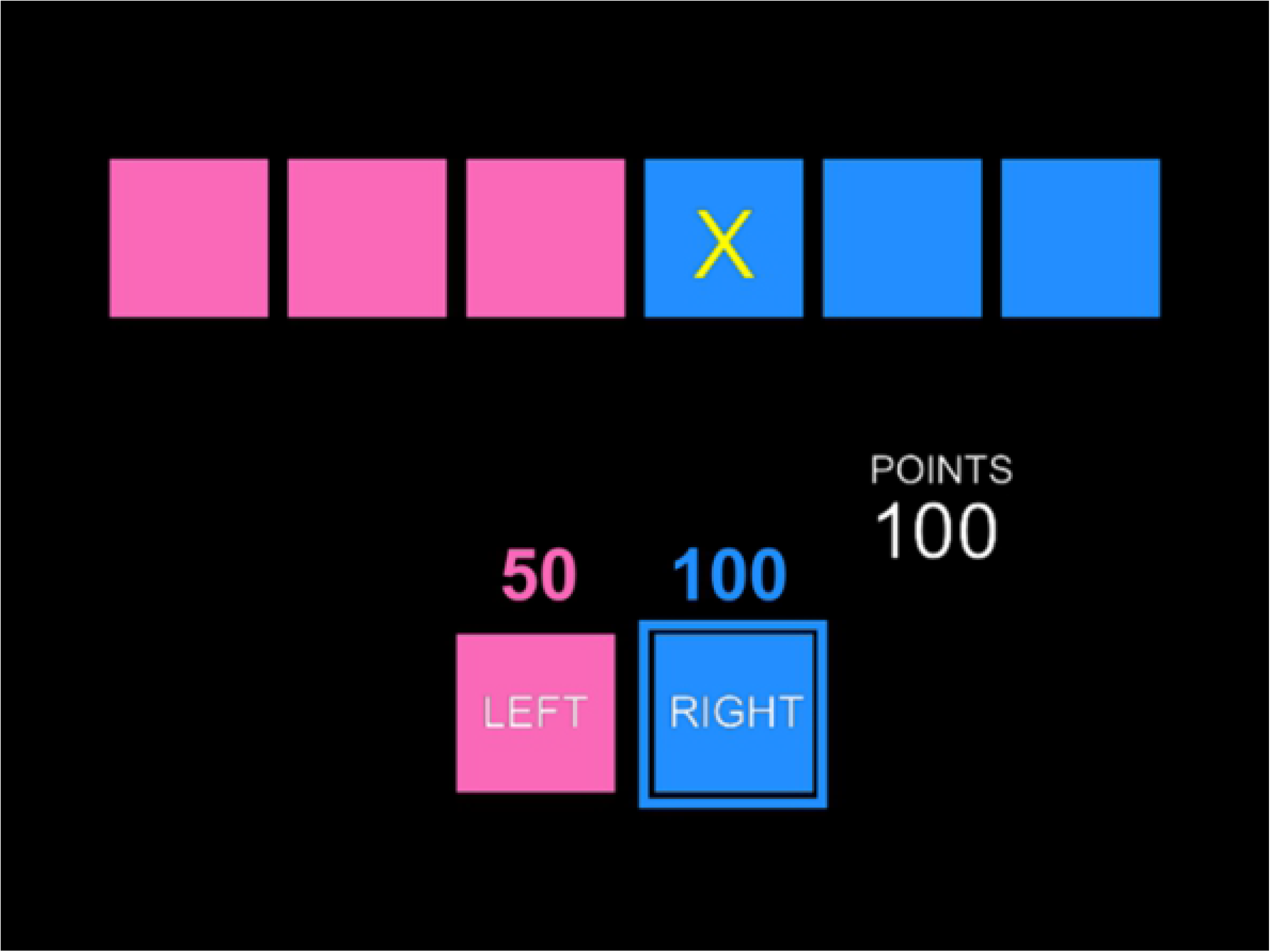
Maastricht Gambling Task. Example screen presenting a trial where choosing blue offers a payout of 100 points with a probability of 3/6 (EV= 50) and pink offers a payout of 50 points and a probability of 3/6 (EV=25). In this example the participant chose the highlighted option (blue) and the token (yellow x) is revealed to be hidden behind one of the blue boxes. In this example the participant gained 100 points as it is presented in white.

Each color has a different bet value representing the potential earnings if the chosen color is correct (hit). A wrong guess results in zero payoff. For example, in Fig 2, the trial offers a chance of 3/6 (50%) of earning 50 points if pink is chosen, and 3/6 (50%) of earning 100 points if blue is chosen. The bet values were selected randomly among five different values (5, 25, 50, 75 or 100) for each color in each trial independently. The participant’s goal was to obtain the maximum of points in each trial. To remove the impact of loss aversion, the MGT does not allow for losses. Trials have no inter-dependency in the MGT. Payoffs are calculated for each trial independently, and are not cumulated over trials. This avoids memory and wealth effects.

Finally, participants see all the possible combinations of the five different bet values with the different probabilities, resulting in 125 unique trials, therefore participants can perceive that there is no deception and all possibilities are randomly assigned. Each trial is displayed twice, which yields a total of 250 trials in random order to guarantee consistent results. Participants had an average of consistency of 100% in the probabilities chosen and 93% (standard deviation 4%, median 93%) in terms of risk-taking and average value chosen across the repeated trials. No participant made significantly inconsistent decisions comparing the two repetitions of each trial in a session.

The token’s location, color distribution and bet values are determined independently and randomly across trials. With this, we guaranteed that there was no deception and full randomization. This also minimizes the chance of any specific strategy development. All participants were informed explicitly that there was no winning strategy since all results were random.

When a participant chooses a color, the choice is highlighted and the position of the token is revealed (Fig 2). Therefore, in this same example, as the participant chose blue and the token was hidden behind a blue box, the participant gets 100 points (as indicated in the white text on the right).

To gain more insight on the different types of trials, we divided them into three clusters according to the differences (or contrasts) in expected values offered by the two options (Pink and Blue), which could capture the difficulty of making a choice in the trial. The lower this difference the more difficult it is for a subject to make a choice. This led to the division of trials in these clusters: low, medium and high contrast. In our analysis, we excluded trials with no difference in expected value, since this group of options includes less trials than the remaining clusters and would not allow balanced analyses. Trials with one strictly dominant option, meaning (for simplicity) trials where the options have differences in expected value > |65| were excluded. This exclusion was made since these were considered non-informative, because these choices are considered obvious and would hardly be affected by any environmental or intrinsic factor. In total 204 out of 250 trials were analyzed per session. The cluster division can be seen in detail in the S1 Table.

### Transcranial alternating current stimulation

We aimed at stimulating the left DLPFC. A small circular (diameter: 2.1cm, thickness: 2mm) and a large (Outer diameter: 11cm; Inner diameter: 9cm, thickness: 2mm) rubber ring tACS electrode (neuroConn, Ilmenau, Germany) were placed using conductive gel (Ten20 conductive Neurodiagnostic electrode paste, WEAVER and company, Aurora CO, USA) onto the left DLPFC, with the small electrode positioned over F3 (based on the international 10-20 EEG system) and the large electrode around it. Electrode positioning and tACS simulation were modelled with SimNIBS (25), as shown in Fig 3.

**Fig 3.**
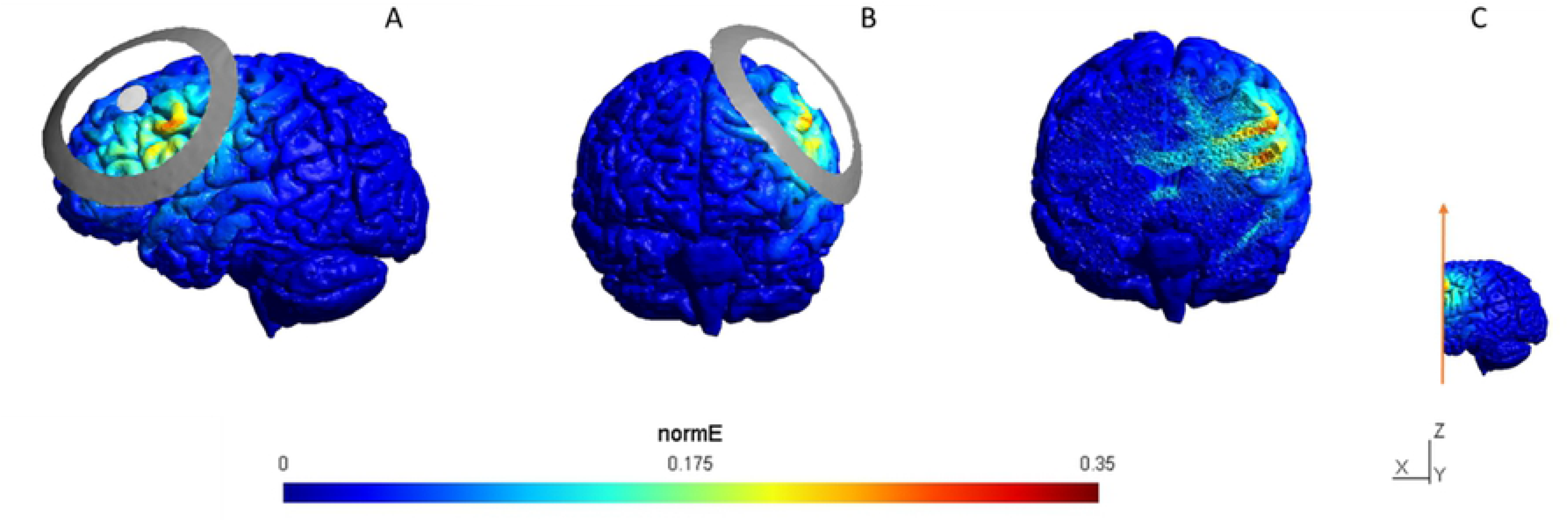
SimNIBS tACS simulation. Left lateral (A), frontal (B) view and of the stimulation and coronal cut at F3 to show the potential subcortical reach of the stimulation (C). Colors stand for the normalized electric field [0-0.35], meaning that the red areas are the areas where the electric stimulation has a higher incidence.

This ring electrode montage enables a higher spatial focality as compared to standard rectangular electrodes (26). Alternating current was applied using a neuroConn DC-stimulator with remote triggering (neuroConn, Ilmenau, Germany) and DataStreamer software (27), for which we created stimulus protocols on Matlab2018b (The Mathworks Inc., Massachusetts, USA) for each condition. Stimulation frequency and intensity were set to 6,5 Hz (theta-range stimulation) and 40 Hz (gamma-range stimulation) and a stimulation intensity of 1,5mA peak to peak, phase offset was set to 0 and 100 cycles were used for ramping up. For sham tACS, the current was ramped up at a 6.5 Hz frequency for 30 seconds and ramped down immediately after. The impedance of the tACS electrodes was kept below 15 kΩ during stimulation. The average stimulation time lasted 30 minutes. Participants were blind to the stimulation protocol and the experimental hypotheses. Questionnaires applied after the experimental session confirmed that participants were unaware of the stimulation protocol.

### Electroencephalography

EEG electrodes were positioned according to the 10-20 international EEG system around the stimulation site (F1 and F5), contralateral to the stimulation site (F2 and F6) and on the parietal cortex (P5 and P6), with Cz being used as reference and the left mastoid used as ground. EEG measurements were done immediately before and after the tACS stimulation, each lasting 3 minutes, to measure resting-state theta-band activity (measurement before the stimulation) and the effects of the entrainment (after stimulation). Participants were asked to stay with their eyes closed, relaxed and to avoid any movement.

Data was recorded (DC-200 Hz, sampling rate 500 Hz) with a BrainAmp Standard EEG amplifier and the BrainVision Recorder software (BrainProducts GmbH, Munich, Germany). Impedance levels were kept below 15 kΩ. Offline preprocessing was conducted using the Fieldtrip toolbox (28) and custom Matlab scripts. EEG recordings were low pass-filtered in the analog domain (cutoff frequency: 250 Hz) and then digitized (sampling rate: 1000 Hz). Offline preprocessing was performed with a notch-filter (50 Hz) to remove electrical noise and demean the data over the full dataset. After that, it was segmented into 90 trials of 2 seconds each. Trials with high variance and excessive noise were excluded by visual inspection and variance analyses.

### Statistical Methodology

We analyzed four different behavioral dependent variables: 1) Risk, 2) Probability scores 3) Value and 4) Response time.

### Risk

The measure of risk-taking behavior should be dependent on both the probabilities of outcomes and the value associated with each outcome. In our experiment, betting on a color *X* (*X* = blue or pink) in a trial with probability *p*, and a payoff of *x* would have an expected payoff of *xp*. For instance, when choosing pink, the probability of being correct (a hit) and getting the reward is equal to the proportion of pink boxes during that trial, and the probability of being incorrect and getting no reward is equal to the proportion of blue boxes. Therefore, the expected payoff from choosing color *X* in a trial is given by the following.

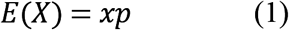

For example, in a trial with 1 blue box with a bet value of 100, and 5 pink boxes with a bet value of 5, the expected payoff for blue and pink are respectively 16.67 and 4.17. This makes blue more attractive for a risk-neutral participant. Therefore, an option is strictly dominant for a risk-neutral participant if it has higher expected payoff.

The measure of risk takes into account the level of variation (29). The variance of payoffs from choosing color *X* is given by the following.

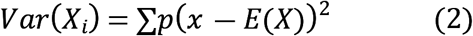

For example, in a trial with 1 blue box with a bet value of 25, and 5 pink boxes with a bet value of 5, the expected payoffs of both options are the same, 4.17. However, the variance of blue (86.81) is much higher than that of pink (3.47). Therefore, the option blue is considered riskier than pink. Therefore, for two bets with the same *expected value*, the one with a larger variance is considered riskier. From variance, we calculated standard deviation (SD) as our measure of risk-taking behavior (e.g. Myerson, 2005) which is our main dependent variable, from now on referred to as Risk.

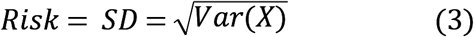

### Probability scores

Previous studies only considered the choice of specific outcome probabilities as an indicator of risk (4,22,31), meaning that in these studies, a choice is typically considered risky if the probability is below 50% and safe if its probability is above 50%. To allow a more refined analysis of participant’s preferences of probabilities, they were transformed into a scale ranging from −2 to 2. The choice of a higher probability was classified with a negative score and that of a lower probabilities received a positive score. In simple terms, these scores indicate that options with a higher level of uncertainty have positive scores, while safer options have negative scores. These probability scores can be seen in Table 1.

**Table 1.** Probability scores. Higher scores indicate that participants chose the trials with lower probabilities (risk prone), while lower scores indicate that participants chose higher probabilities (risk averse). For example, if a participant choose Blue in a trial where the distribution of Blue boxes is 1/6 (and Pink boxes 5/6) the participant would receive a score 2, indicating that the participant chose the lowest probability possible. If in this same trial the participant chooses Pink, the score would be −2, indicating that this participant chose the highest possible probability.

| Pink | Blue | Choice | Probability |
| --- | --- | --- | --- |
| 5 | 1 | Blue | 2 |
| 1 | 5 | Pink | 2 |
| 4 | 2 | Blue | 1 |
| 2 | 4 | Pink | 1 |
| 3 | 3 | Pink | 0 |
| 3 | 3 | Blue | 0 |
| 4 | 2 | Pink | -1 |
| 2 | 4 | Blue | -1 |
| 5 | 1 | Pink | -2 |
| 1 | 5 | Blue | -2 |

### Value and Response Time

To analyze the average value chosen by a participant in each session, their choices of bet values independent of the trial result (being correct or incorrect) were averaged. That variable is named Value. Furthermore, response times (RT) were also recorded for every decision.

### Behavioral data analyses

The behavioral data were preprocessed using custom Matlab (The Mathworks Inc., Massachusetts, USA). We performed a series of Linear Mixed Model analyses to estimate the effects of stimulation (sham, theta and gamma) on risk-taking behavior. Our final models were fixed effects models, with participant-specific random effects. All the analyses presented normally distributed residuals, showed no heteroscedasticity, and no observations were removed as outliers.

Overall, we constructed linear mixed models where each observation is a unique subject-cell pair. Each cell is a unique combination of session and contrast. That is, 3 sessions by 3 levels clusters (LC (low contrast), MC (medium contrast) and HC (high contrast)), resulting in 9 unique observations (cells) per subject. The resulting models can be represented as the following:

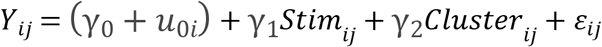

*Y*_*ij*_ stands for each of the behavioral outcome variables; *i* stands for the the i-th participant and *j* represents the j-th cell; γ_0_ stands for fixed effect intercept; *u*_0i_ stands for the subject-specific random effect; *Stim* stands for Stimulation condition (sham, theta, gamma), *Cluster* stands for the three different levels of contrast of the trials (low, medium, high contrast). Stim and Cluster are subject-cell specific, hence the subscript *ij*. We checked correlation among the behavioral dependent variables, and checked the robustness of our results for each behavioral dependent variable when using appropriate controls of other behavioral outcomes. This did not affect our main results.. Results were confirmed with additional repeated measures ANOVA analyses omitted for conciseness.

For the analysis of Response Time (RT), we compared covariance structures on Maximum Likelihood (ML) estimations using AIC. For these analyses, the best fit was using a Heterogeneous Toeplitz (TPH) covariance structure.

In the analyses of Value, Probability scores and Risk, the covariance structures were compared on Reduced Maximum Likelihood (REML) estimations using AIC. The covariance structure used for Value and Risk were Heterogeneous Compound Symmetry (CSH). The final model to analyze Probability scores used a TPH covariance structure for repeated measures.

### EEG analyses

We preprocessed the data separately for low (1-20 Hz) and high (20-90 Hz) frequencies. For low frequencies, a fast Fourier transformation was performed with hanning tapers and output frequencies between 1 and 20 Hz. For high frequencies, a fast Fourier transformation was performed with discrete prolate spheroidal sequences (DPSS) tapers, smoothing factor of 5 Hz and output frequencies between 20 and 90 Hz. Then data were Log normalized to control for discrepancies driven by individual variability (32).

To look for differences in theta and gamma power before and after the stimulation protocols, the power spectra were averaged for the before and post stimulation measurements. Theta-band was defined between 5 and 8 Hz, with 1.5 Hz above and 1.5 Hz below the stimulation frequency (6.5 Hz), whereas gamma-band was defined between 35 and 90 Hz. Theta and gamma power were analyzed for all channels pre and post stimulation, with focus on the frontal left channels (F1 and F5) around the stimulation focus, frontal right channels (F2 and F6) contralateral to the stimulation.

To investigate whether a change in the hemispheric relationship in theta power took place, we calculated the average of theta power in the right hemisphere minus the average in the left hemisphere, named frontal asymmetry (right - left) (9). Moreover, we compared the changes in theta as well as gamma power in the parietal channels before and after stimulation to analyze how focal were the stimulation effects.

The effects of stimulation in each condition were compared within participants for the intervals of 1 minute and three minutes. Signal processing and EEG data preprocessing were conducted using Matlab (The Mathworks Inc., Massachusetts, USA)custom scripts and the Fieldtrip toolbox (28). The difference in theta power across conditions was correlated with the behavioral results both using the theta-asymmetry before stimulation as covariate and the changes in theta and gamma frequencies as dependent variables by performing a repeated measures ANCOVA with Bonferroni correction.

## Results

### Behavioral results

#### Main results: Risk

To analyze the effects of stimulation on risk-taking behavior, measured as the average standard deviation of the chosen option (as described above), we ran a Linear Mixed Model analysis using stimulation protocol as factor. The estimated fixed effects showed a significant reduction of −0.301 on risk-taking behavior during theta-band stimulation, t(66.693)=−2.035, p=.046, SE= .148, compared to sham (Fig 4). Moreover, gamma stimulation did not affect the participant’s average risk-taking significantly compared to sham, t(69.992)=−1.224, p=.225, SE= .100, confirming that the effects observed are frequency specific.

**Fig 4.**
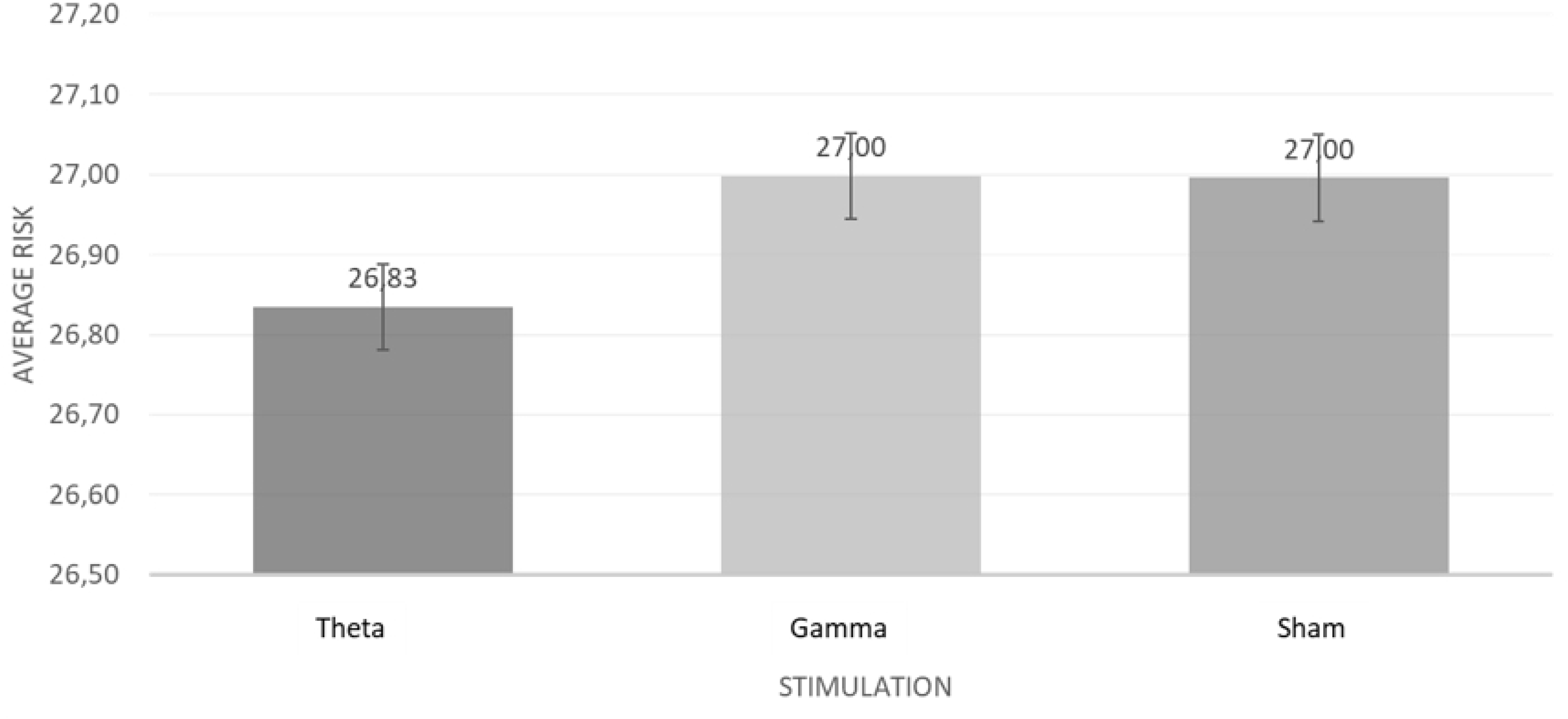
Average Risk. Estimated by the average standard deviation of each participants’ choice across stimulation conditions (Theta [6,5 Hz], Gamma [40 Hz] and Sham). Error bars depict SEM. Risk can vary between 11.750 and 36.150.

#### Probability scores

The Linear Mixed Model analyses with Probability as the dependent variable did not yield significant main effects for stimulation, F(2,47.287)= .762, p=.921. The estimated fixed effects analyses did also not yield significant effects of theta-band stimulation, t(35.078)=.316, p=.754, SE= .014, or gamma stimulation, t(67.703)=−.801, p=.426, SE= .009, when compared to sham, meaning that no significant differences in the probability scores were observed after the different stimulation protocols.

#### Value

The analyses of the effect of the stimulation on the average values were run using a Linear Mixed Model Procedure with Value as the dependent variable, which yielded a non-significant main effect of stimulation, F(2,91.890)=2.427, p=.094. Further analyses of estimated fixed effects yielded significant effects of Theta stimulation on value, with a reduction of −0.674 compared to sham, t (64.331)=−2.130, p=.037, SE= .316. No significant effects were observed after Gamma stimulation t(71.190)=−1.270, p=.208, SE= .207, compared to sham. This means that there was a significant reduction in the average value chosen by the participants due to the Theta stimulation, confirming that this is a frequency exclusive effect (Fig 5). These findings reinforce the strong relationship between risk-taking behavior and valuation since both processes were affected by the same pattern of stimulation.

**Fig 5.**
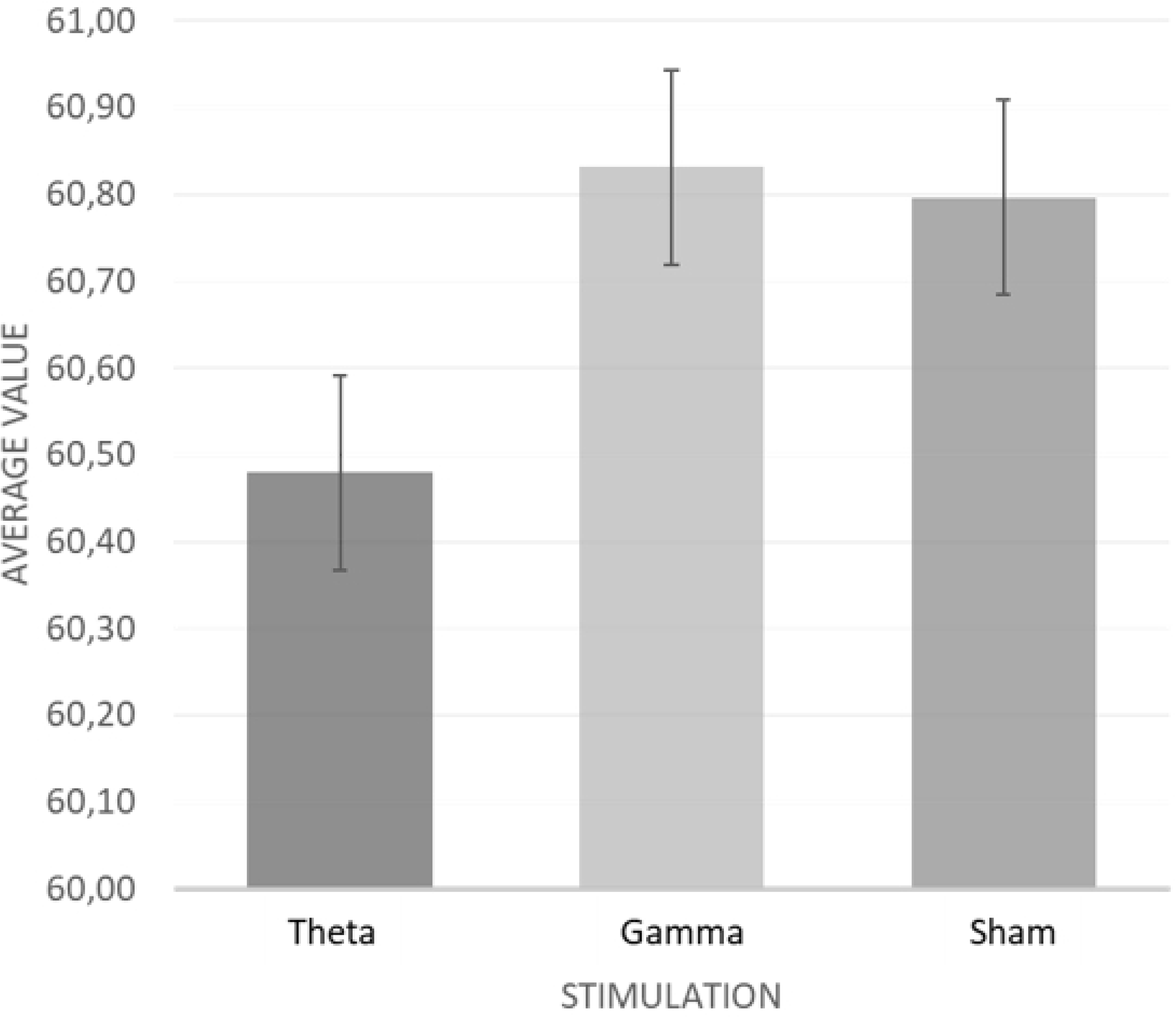
Average Value. Average value selected by the participant in each condition (Error bars depict SEM).

#### Response Time

To analyze the participant’s response time (RT), we used a Linear Mixed Model with RT as dependent variable. The results show strong significant effects of stimulation, F(2,50.240) =35.803, p<.001.

Estimated fixed effects analyses yielded significant results for theta stimulation, t(24.259) =5.161, p<.001, SE= .066 and nearly significant effects for gamma stimulation, t(63.862)=1.880, p=.065, SE= .027, when compared to sham. Theta stimulation led to an increase of 41.11% in response time (compared to sham). This implies that the theta stimulation led to an increase in the deliberation time, which cannot be attributed to the stimulation per se, since this effect was only marginally significant in the gamma stimulation condition. Details can be observed in Fig 6, where response time is plotted against contrast, or trial difficulty level based on the cluster division previously explained, from easier decisions (which are clear, with big differences in EV between Pink and Blue) to difficult decisions in which the mental calculation to define the most advantageous option is more challenging.

**Fig 6.**
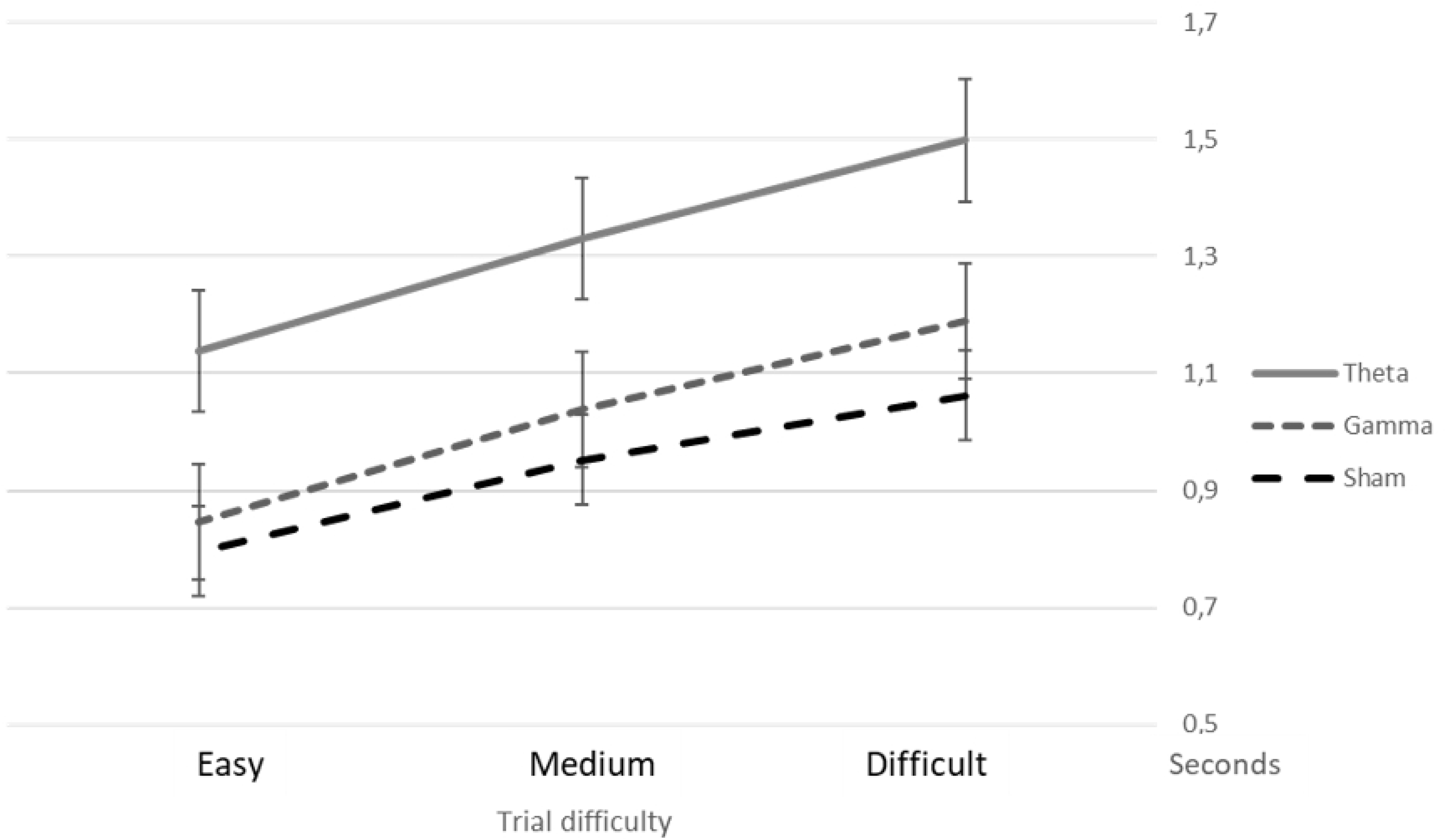
Average Response Time. Presented in seconds by trial difficulty (easier trials have a higher difference in expected value between the trials) by Stimulation protocol (Error bars depict SEM).

### EEG results

#### Theta-band entrainment

To investigate the effects of theta-band stimulation on EEG results, we ran a repeated measures ANOVA with theta power as dependent variable. The repeated Measures ANOVA used a 3 (stimulation condition: theta, gamma and sham) by 2 (time: before and after stimulation) by 6 (theta power averaged over 3 minutes on each electrode: F1, F5, F2, F6, P5 and P6) within subject design, with Bonferroni correction for multiple comparisons.

The results showed a significant main effect of time, with theta power increasing from an average of −0.135 to an average of −0.056 after stimulation, F(1,6) =3.383, p=.009. There was no significant main effect of stimulation, F(1,12) =.824, p=.627, and no significant interaction effect between stimulation and time, F(1,12) =.822, p=.629 (for descriptives please refer to S2 Table).

Further analyses included a 3 (stimulation condition) by 2 (time) repeated measures ANOVA using frontal asymmetry as dependent variable. There was no significant effect of stimulation on frontal asymmetry, F(1,2) =1.191, p=.168; time, F(1,1) =.059, p=.810 or of the interaction between time and stimulation, F(1,2) =.807, p=.457.

*Post hoc* analyses were done to investigate whether the effect might be more robust during the first minute immediately after stimulation. To do so we ran a repeated measures ANOVA using as dependent variable the difference in theta power between the first minute before stimulation and the first minute immediately after it. We used a 3 (stimulation conditions: theta, gamma and sham) by 7 (theta power difference on each electrode: F1, F5, F2, F6, P5, P6 and change in frontal asymmetry) within subject design, with Bonferroni correction for multiple comparisons. These analyses yielded a significant effect of stimulation, F(1,12) =4.440, p<.001.

Further contrasts showed significant effects of both theta and gamma-band stimulation. Of interest, there was a significant effect of theta-band stimulation (and not gamma) on asymmetry change (pre-post) when compared to sham t(2)= 2,528, p=.012. However, this effect is mainly driven by a decrease in asymmetry in the sham condition, which could be observed in Fig 7, indicating that the decrease is due to the task execution and not due to the stimulation.

**Fig 7.**
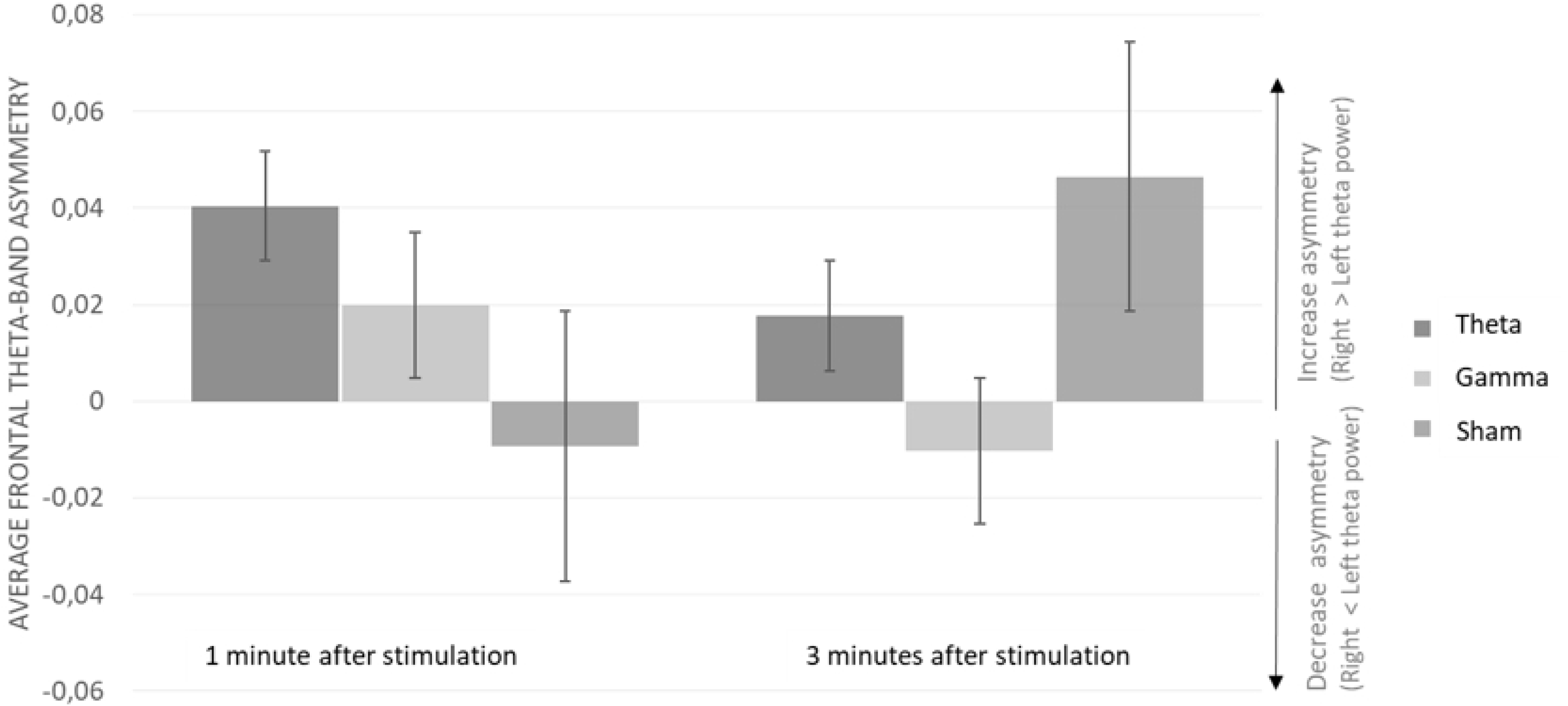
Frontal theta-band asymmetry difference during 1 and 3 minutes after stimulation. Average theta power change [5-8 Hz] (pre-post) per stimulation protocol. Graph shows average data collected during 1 minute after stimulation (left) and 3 minutes after stimulation (right).

According to our analyses, the asymmetry went from a negative score in the first minute after the task to a positive score when 3 minutes are analyzed. This indicates that during the first minute after the task execution there was a higher theta power in the left hemisphere, which changed during the following minutes, since the analyses including 3 minutes after task indicate a higher theta power in the right hemisphere. More details can be seen in Fig 7. Moreover, the only frequency specific effect observed after theta-band stimulation was during the first minute post stimulation in F6 (contralateral to the stimulation focus) (t=3.876, p<.001).

We also conducted a partial correlation analysis between the frontal theta asymmetry, theta power in F1, F3, F5, F6, P2 and P6 and the behavioral responses (probabilities chosen, average value chosen, risk and response time). The level of asymmetry before or after the stimulation did not significantly correlate with either of the behavioral measures. Theta power in F1 and F2 were significantly correlated to the probabilities chosen (r= .107, p=.018 and r= .013 and p=.013 respectively), although there were no significant effects of stimulation on the probabilities chosen by the participants. The results also indicate trends regarding the correlations between theta-power in F1 and F2 and the average values chosen (r= .075, p=.098 and r =.077 and p=.090 respectively) and between theta-power in these same electrodes and risk (r=.080, p=.077 and r=.082, p=.071 respectively).

The inclusion of asymmetry of theta-power in any of the electrodes in the regression models used to analyze the behavioral results did not improve the fit of these models and therefore was discarded.

#### Gamma-band entrainment

The effects of gamma-band stimulation were investigated using a 3 (stimulation condition: theta, gamma and sham) by 2 (time: before and after stimulation) by 6 (theta power averaged over 3 minutes on each electrode: F1, F5, F2, F6, P5 and P6) within subject repeated measures ANOVA, with Bonferroni correction for multiple comparisons. No significant effect of stimulation condition F(1,12) =0.845, p=.604, nor of time, F(1,6)=1.045, p=.424 or the interaction between stimulation and time, F(1,12) =1.035, p=.465 was observed,

## Discussion

The present study aimed at investigating the functional relationship between frontal theta-band oscillations and risk-taking behavior. Although previous studies (9,10) have shown a correlation between resting state frontal theta band asymmetry and risk-taking behavior, no direct causal relationship has thus far been shown. We hypothesized that theta oscillations underlie the neuronal communication for recruiting DLPFC when the decision-making process includes risk, being fundamental for the modulation of risk-taking behavior (12). We therefore expected theta-band stimulation to cause a reduction in risk-taking behavior and that this effect is frequency specific.

As predicted, we were able to effectively reduce risk-taking behavior in healthy participants using theta-band tACS over the left DLPFC, compared to sham and gamma-band stimulation. These findings confirm the functional relationship between theta-band frequencies and risk-taking behavior regulation, being a fundamental part of the electrophysiological mechanism responsible for this modulation. Theta band tACS leads to a significant decrease of 1.12% of risk taking behavior compared to sham. This was not the case during gamma stimulation. To our knowledge, our study is the first to show the frequency specificity of this effect. Moreover, we observed a significant reduction on value sensitivity due to theta-band (and not gamma) stimulation, meaning that participants opted for lower values after theta-band stimulation compared to the results obtained in the sham or gamma conditions. These results are in line with previous studies, where participants became more risk-averse after non-invasive brain stimulation with reduced sensitivity to value (22,33–35). However, our study was able to show that also this effect is frequency specific. Therefore, it is expected that theta-frequencies would play a fundamental role in the reduction of value sensitivity, meaning the recruitment of DLPFC as executive control to modulate the VMPFC response to the value (36).

The stimulation did not affect the probabilities chosen by the participants, indicating that the choice of probabilities might be regulated by a different electrophysiological mechanism. Even though our results indicate that probabilities and value are evaluated independently in our brain, behaviorally and in terms of neurological activity these processes are at least strongly correlated (2,29,37). This means that both inputs are considered (bet value and its probabilities) in order to inform the decision process, which justifies the use of standard deviation as an estimation of risk. Our approach considers the option’s expected value (meaning the bet’s probabilities and value) to estimate risk, which is in our perspective a more naturalistic evaluation of risk. Our findings indicate that participant’s reductions on risk-taking behavior were mainly driven by a reduction on the average value sensitivity.

Although we did not have a specific hypothesis regarding the response time, it is interesting to notice that theta stimulation increased response time compared to sham and gamma stimulation. It may be speculated that the increased response time reflects a longer deliberation process (38).

It is important to note that our results contradict the study by Sela and colleagues (2012).Their results indicated an increase in risk-taking behavior after theta-band stimulation, which might be explained by their choice of the Balloon Analog Task as experimental paradigm. Since this task has a strong factor of impulsivity the effect observed should reflect an increase in impulsivity and not in risk-taking behavior (16). Moreover, they considered the tolerability to losses (measured as sequential explosions) as an indicator of riskier choices (14), which means that their results might also indicate a reduction in loss-aversion. Since our experimental paradigm (MGT) avoids loss-aversion and impulsivity, we may have more directly assessed risk-taking behavior.

In addition to assessing the behavioral effects of our oscillatory brain state neuromodulation on risk taking modulation, we also used EEG to measure oscillatory activity before and after the tACS stimulation. When comparing theta power before and after theta tACS, no significant changes were found, nor did we reveal significant changes in hemispheric theta-band asymmetry after theta-band stimulation. This may seem surprising and in contrast to our behavioral effects being attributed to and interpreted as being caused by tACS-induced increased in left theta power. However, it is important to note here that while behavioral effects were assessed during tACS being applied simultaneously with task execution, the EEG measurements, due to tACS artefacts, were restricted to assessing the oscillatory activity after both the behavioral performance and the tACS stimulation had ended. Especially the latter may be a straightforward explanation for the absence of significant EEG effects in a pre-post tACS design as such effects rely on a significant longer lasting neurophysiological effect of tACS beyond the period of stimulation itself.

However, the question whether or not tACS-induced entrainment is longer lasting is far from being settled (39). Various previous studies also reported difficulties in establishing longer lasting effects of tACS on excitability or neural plasticity (13,40–42). Considering our results, we may therefore speculate that the EEG effects were only present during the task and stimulation and faded away immediately after tACS had ended. Our post hoc analyses focusing only on the first minute of post EEG measurements after TACS and contrasting these effects to the entire post EEG period indicate time-sensitive changes in theta band asymmetry in line with this speculation. Yet, our study was not designed to conclusively test and other related hypotheses regarding the difference between the immediate versus lasting effects of tACS on neural oscillatory activity. Follow up studies with online measurements using algorithms to remove stimulation artifacts could be used to investigate such possibility, although currently this methodology is still in debate (42).

In addition, we also revealed that the task execution itself had lasting effects on theta band asymmetry as indicated by *post hoc* analyses of the EEG measures immediately after the task execution in the sham condition. In other words, unrelated to tACS, the mere behavioral performance in risk taking modulation tasks considerably affected theta band asymmetry after task execution had been completed. At the same time, our behavioral results showed no significant correlation with resting-state frontal theta-band asymmetry at baseline, indicating that these effects cannot be explained only by the resting-state frontal asymmetry or by changes in asymmetry due to the stimulation.

The stimulation frequency-specificity of our significant behavioral findings, however, confirming our a priori hypothesis that specifically theta, not gamma or sham, neurostimulation should affect risk taking behavior, clearly represents supporting evidence for the functional relationship between theta-band stimulation and risk-taking regulation driven by reduction of sensibility to reward. This work also contributes to the understanding of the frontal areas interaction in the regulation of risk-taking behavior, as much as the role of theta-band oscillations in this process. Moreover, it gives insights about the causes of individual differences in risk-taking, granting the analysis of frontal resting state brain activity a potential role to infer differences in individual risk-proneness. This can be used in the construction of more accurate economic models of risk-taking, or contribute to the development of diagnosis and intervention techniques for patients with abnormal risk-taking behavior, since this is characteristic of a range of psychiatric and neurological disorders (43).

## Conclusion

Although it is widely accepted that DLPFC have an important role in risk-taking regulation it is not clear how the recruitment of this area occurs in presence of risk. Theta oscillations are potentially responsible for neuron communication when cognitive control is needed (12). In our study, we provided empirical evidence for the direct functional relationship between prefrontal theta band activity and risk-taking regulation using high definition theta-band tACS with gamma-band entrainment and sham as control. A significant reduction of risk-taking behavior was observed after theta-band, but not gamma-band or sham tACS over left DLPFC, confirming the specific role of theta frequencies in risk-taking behavior regulation. Such findings indicate that prefrontal theta-band oscillations are potentially the basis of communication between frontal areas during risk-taking regulation.

## Acknowledgements

We would like to thank our research assistant Kira Temme for her help during data collection.

